# Deciphering Precursor Cell Dynamics in Esophageal Preneoplasia via Genetic Barcoding and Single-Cell Transcriptomics

**DOI:** 10.1101/2025.02.26.637920

**Authors:** Jinho Jang, Kyung-Pil Ko, Jie Zhang, Sohee Jun, Jae-Il Park

## Abstract

Although histologically normal, esophageal preneoplastic cells harbor early genetic alterations and likely exhibit lineage plasticity. However, their origins and trajectories remain unclear. To address this, we combined genetic barcoding with single-cell RNA sequencing to trace the lineage of esophageal preneoplastic cells. We identified a distinct progenitor-like cell population with high plasticity. Through a newly developed scoring system, these high-plasticity cells are mapped, revealing their contributions to proliferative and basal cell populations. This approach uncovers molecular markers, including *Nfib* and *Qk*, that define these precursor cells, validated by spatial transcriptomics and a *Trp53 Cdkn2a Notch1* mouse model. These findings provide critical insights into early tumorigenesis, highlighting the potential of precursor cells as biomarkers for early detection and therapeutic targets of esophageal squamous cell cancer. By elucidating the cellular dynamics underlying esophageal preneoplasia, this research lays the foundation for strategies to prevent malignant progression, offering broader implications for improving cancer diagnostics and treatment approaches.

**Significance Statement:** Preneoplastic cells often appear histologically normal yet carry early genetic and transcriptional changes that predispose them to malignant transformation. In this study, we combine genetic barcoding with single-cell transcriptomics to uncover the lineage dynamics of esophageal preneoplastic cells. We identify a distinct progenitor population, preneoplastic cells of esophageal squamous cell carcinoma (pESCC), characterized by high plasticity and a unique trajectory that gives rise to proliferating and basal cell populations. By developing a new computational scoring method to integrate lineage topology with differentiation state, we provide a framework for tracing cellular origins beyond conventional inference-based models. Our findings shed light on the earliest events in tumor initiation and offer a new paradigm for identifying biomarkers and intervention targets in the precancerous stages of esophageal cancer.

---

Cancer cells exhibit extensive heterogeneity arising from diverse cell lineages, which contributes to drug resistance and tumor relapse (1, 2). Understanding these cell lineages has long been a major challenge in understanding the biology of cancer and establishing effective therapeutic strategies. However, despite extensive studies of cancer cells, the identification of cancer stem cells and the related cell lineages remains elusive (3, 4). This challenge is exacerbated by the clonal diversity within neoplastic cells, which intensifies cellular heterogeneity and complicates lineage relationships. Since intra-tumor heterogeneity arises from both genetic and non-genetic mechanisms from the early phase of tumor development, therapeutic strategies targeting genes or cells in late-stage tumors often fail to achieve durable tumor control (5). Therefore, targeting core cell lineages during the early stages of tumor development, before clonal expansion, is crucial. Recent studies have also reported that preneoplastic cells acquire cell plasticity, granting them the potential to transition into neoplastic cells (6-8). Elucidating the unique high-plasticity cell populations and lineage trajectories of preneoplastic cells, which are distinct from normal cells, could provide critical insights for preventing cancer development.

Experimental approaches used to identify cells of origin have included genetic reporter and barcoding systems (9-11). Genetic reporters, such as those encoding fluorescent proteins or beta-galactosidase, have been widely used for permanent labeling of parent cells through spatiotemporal genetic recombination using Cre-loxP or Flp-Frt systems. When controlled by tissue- or cell-specific promoters, these genetic recombination systems enable precise tracking of cellular lineage (12). Despite their broad applications for in vivo models, genetic reporter systems typically rely on a limited set of researcher-selected genes for cell labeling, which introduces potential bias (13, 14). Moreover, the ability to trace multiple cell lineages is constrained by the limited number of available fluorescent tags with non-overlapping emission spectra (11). Although stochastic and combinatorial expression models for fluorescent proteins have recently been introduced, the number of distinguishable colors remains restricted, and the persistence of fluorescence can present additional challenges (15, 16). Another significant limitation is the ubiquitous expression of reporters across all cells within the same lineage. Thus, the genetic reporter-based approach captures only a single snapshot of lineage dynamics at a specific time point, making the interpretation of lineage directionality more subjective and potentially less reliable.

Genetic barcoding has also been used to track large numbers of cells by integrating diverse barcode libraries into DNA, which are subsequently inherited by daughter cells originating from stem cells (17, 18). With advancements in next-generation sequencing, this technique has been successfully used to map cellular differentiation trajectories from stem cells (19). Meanwhile, single-cell RNA sequencing (scRNA-seq) has addressed several limitations of earlier methods by enabling transcriptomic analysis at the single-cell level. Computational tools such as RNA velocity, CytoTRACE, and signaling entropy have been developed to reconstruct cell lineages based on transcriptional dynamics (20-22). Although these tools have helped to reduce biases, they still have inherent limitations, as they rely on a restricted set of prior knowledge and predict cell lineages through computational inference, which may introduce uncertainties.

These significant advances notwithstanding, identifying stem cells and tracing the origins of cancer cells remain formidable challenges. This difficulty primarily arises from the high cellular heterogeneity and plasticity of cancer cells, which can undergo de-differentiation or trans-differentiation, mechanisms that are rarely observed in normal cellular hierarchies (1, 2, 23). These dynamic transitions obscure lineage relationships and complicate the identification of tumor-initiating populations. Therefore, novel tools are needed to accurately resolve these atypical cell states without overreliance on inference-based predictions or prior biological knowledge.

Esophageal tissue consists of relatively simple cell types, and its normal epithelial cell lineage has been well characterized. However, esophageal squamous cell carcinoma (ESCC) exhibits significant heterogeneity, with disrupted and ambiguous cell identities serving as a hallmark of this cancer type (24). Studies of genomic data from patients with ESCC and experimental models have revealed that key genomic alterations occur before the onset of neoplastic tumor growth (25-27). Although these genetic alterations alone are insufficient to drive malignant transformation, cells harboring them display transcriptomic profiles closely resembling those of neoplastic cancer cells, underscoring their critical role as a potential source of cancer development (27).

In this study, we introduce a novel tool to overcome the current limitations of lineage tracing and to identify the cells of origin in preneoplastic lesions. Characterizing preneoplastic cells at an early stage represents a promising strategy for intercepting tumor progression and preventing the transition into fully malignant states. The findings from this study, which focus on identifying the origins of esophageal preneoplastic cells, provide valuable insights into potential strategies for improving the prognosis of patients with esophageal cancer by uncovering the early molecular events of neoplasia.

## Results

### Establishment of cell lineage networks by using iTracer vectors and scRNA-seq analysis

To identify the unique features of preneoplastic cells, we used a genetically engineered mouse esophageal organoid model for preneoplasia, as described previously (27). The combinatorial knockout (KO) of *Trp53* and *Cdkn2a* results in organoids that maintain normal morphology while exhibiting transcriptomic profiles closely resembling those of neoplastic models, indicating a preneoplastic phenotype (27). Briefly, esophageal organoids derived from *Trp53* ^f/f^ mice were infected with an adenoviral vector expressing Cre, followed by transduction with a CRISPR/Cas9 system targeting *Cdkn2a* using single guide RNA, thereby generating *Tr<u>p</u>53 <u>C</u>dkn2a* double KO (PC) organoids (Fig. 1*A*) (28). These PC organoids were subsequently cultured in a 2D system to assess their autonomous proliferative capacity in the absence of growth factor-enriched medium.

**Fig. 1.**
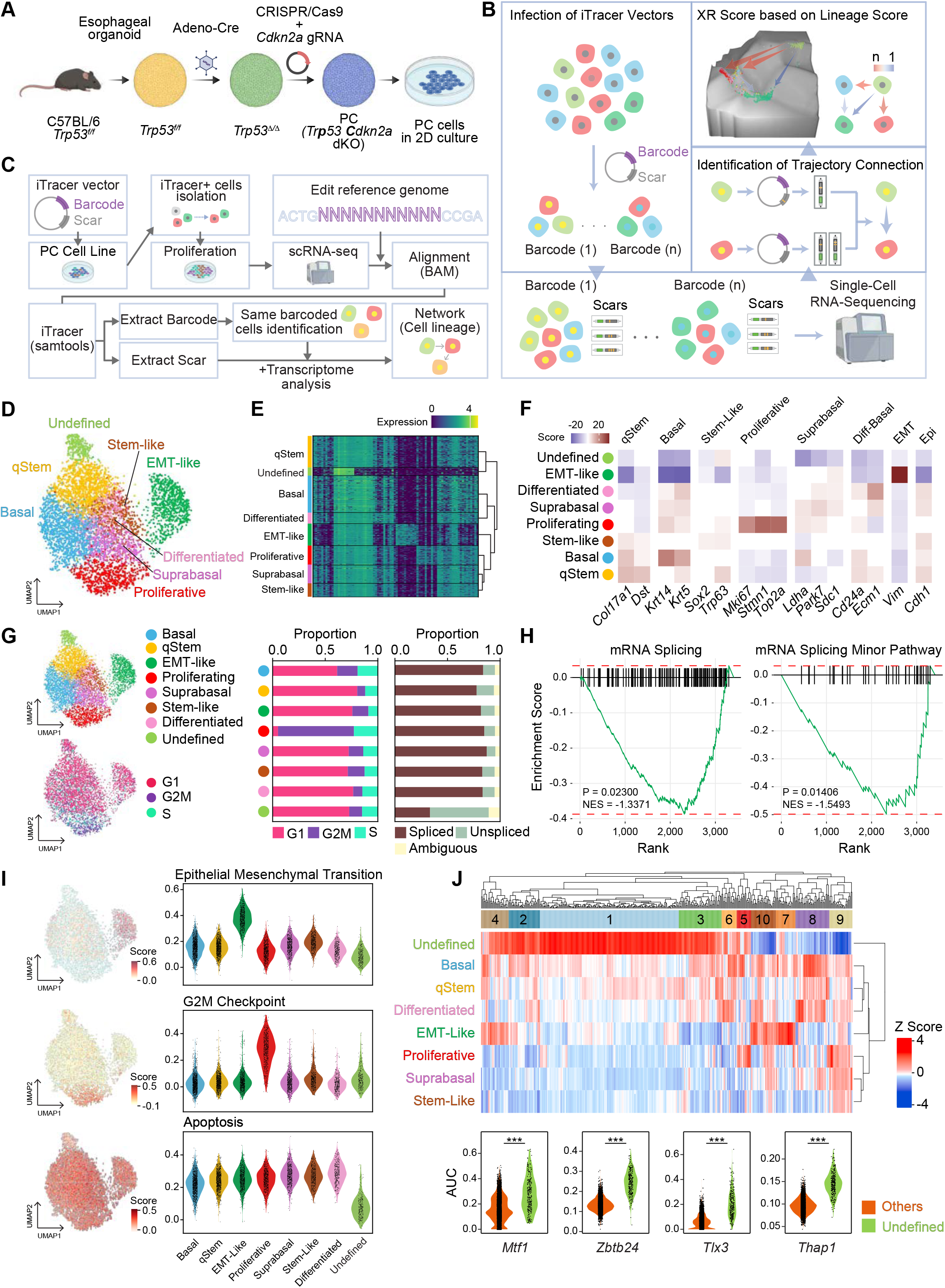
Single-cell lineage tracing and characterization of cellular heterogeneity in esophageal preneoplasia. **(A)** Schematic of the experimental workflow for generating mouse esophageal preneoplastic cells. KO, knockout. **(B)** Schematic representations of analytic pipelines for barcode- and scar-based lineage tracing. **(C)** Detailed workflow of the integrated iTracer and single-cell RNA sequencing analysis pipeline. **(D)** Uniform manifold approximation and projection (UMAP) with annotated cell clusters. EMT, epithelial-to-mesenchymal transition. **(E)** Heatmap showing distinct transcriptional profiles across cell clusters. **(F)** Heatmap displaying key marker gene expression across cell types, with scores from the rank_genes_groups function in Scanpy colored brown (high) and purple (low). **(G)** Cell cycle phase analysis, indicating the distribution of cells in G1, G2/M, and S phases across different clusters. **(H)** Gene set enrichment analysis demonstrating a significant downregulation of mRNA splicing pathways in the undefined cluster (*P* = 0.02300, NES = –1.3371) and minor mRNA splicing pathways (*P* = 0.01406, NES = –1.5493). **(I)** UMAP (left) and violin plot (right) illustrating differential activation of key pathways across cell clusters. **(J)** Heatmap displaying regulon analysis results, highlighting highly activated transcriptional regulators in the undefined cluster. Violin plots visualize the top four enriched factors: *Mtf1, Zbtb24, Tlx3*, and *Thap1* (Student’s *t*-test, \*\*\**P*□<□0.001; 261 cells for undefined and 4,907 cells for others).

To trace the lineage of preneoplastic cells, we used iTracer, a lineage recorder that combines a genetic barcode with CRISPR/Cas9-induced scar systems (29). This system integrates 11-nucleotide random DNA sequences into each cell, which are inherited by daughter cells. Over successive generations, CRISPR/Cas9-mediated genomic insertions or deletions (indels) accumulate, allowing the reconstruction of cellular hierarchies and lineage relationships.

The iTracer plasmid library was prepared as previously described and transfected into 2D-cultured preneoplastic PC cells (Fig. 1*B*) (29). iTracer-introduced PC cells were isolated via flow cytometry and subjected to single-cell RNA sequencing (scRNA-seq). The reference genome was specifically modified to include sequence information for barcode and scar regions, enabling precise lineage tracing through unique genetic markers (Fig. 1*C*). From the transcriptomic profiles, we identified 5,167 out of 17,703 cells exhibiting detectable green fluorescence protein (GFP) transcripts, confirming successful expression of the transduced barcodes and scars (Fig. 1*C*). Barcodes and scars were extracted from the GFP-positive cells by using Samtools, facilitating the construction of lineage networks and enabling a detailed analysis of cellular trajectories and clonal relationships (Fig. 1*C*).

### Single-cell transcriptomics-based dissection of cell clusters in esophageal preneoplasia

We used scRNA-seq and identified seven distinct cell-type clusters based on marker gene expression, with one cluster remaining undefined (Fig. 2*A-C* and *SI Appendix*, Fig. S1*A*). The proliferating cell cluster exhibited significant G2M enrichment, indicating active cell division (Fig. 1*G*), whereas the undefined cluster showed an unexpected accumulation of unspliced transcripts, suggesting an impairment in mRNA splicing (Fig. 1*H*). Gene set enrichment analysis and gene scoring analyses revealed a consistent downregulation of mRNA splicing pathways in the undefined cluster (Fig. 1*H* and *SI Appendix*, Fig. S1*B* and *C*). We also observed upregulation of the G2M checkpoint in the proliferating cluster and increased expression of the epithelial-to-mesenchymal transition (EMT) pathway in the EMT-like cluster (Fig. *1I*). Notably, the undefined cluster exhibited downregulation of apoptosis pathways, ruling out the possibility that it represents a population of dead cells (Fig. 1*I*). Further regulatory network analysis revealed a high activation of multiple regulons in the undefined cluster (Fig. 1*J*), with the top four being *Mtf1, Zbtb24, Tlx3*, and *Thap1*. Among these, *Mtf1* has been reported to promote tumor progression by activating downstream targets (30, 31), whereas *Tlx3* is associated with cell self-renewal and cancer stemness, particularly in prostate cancer (32, 33). The significance of these four TFs as a specific signature for pESCCs was further supported by Receiver Operating Characteristic (ROC) curve analysis (*SI Appendix*, Fig. S1*D*). To examine functional annotations, we performed DAVID analysis on the top 1,000 downstream target genes for each TF, ranked by adjacency scores from pySCENIC. This analysis indicated that the cells are in a state of transcriptional poising (*SI Appendix*, Fig. S1*E*). Specifically, although a core splicing defect leads to functional arrest, the key TFs appear to drive a compensatory response through targets enriched in ‘RNA splicing’ and ‘mRNA processing’ pathways (*SI Appendix*, Fig. S1*E*) (34). At the same time, their target genes are enriched in categories such as ‘regulation of transcription’ and ‘chromatin remodeling’, suggesting involvement in transcriptional reprogramming, and include pathways associated with future growth, such as ‘protein transport’ and ‘cytoskeleton organization’ (*SI Appendix*, Fig. S1*E*) (35-37). These findings suggest that the undefined cell cluster may have a crucial role in tumorigenesis.

**Fig. 2.**
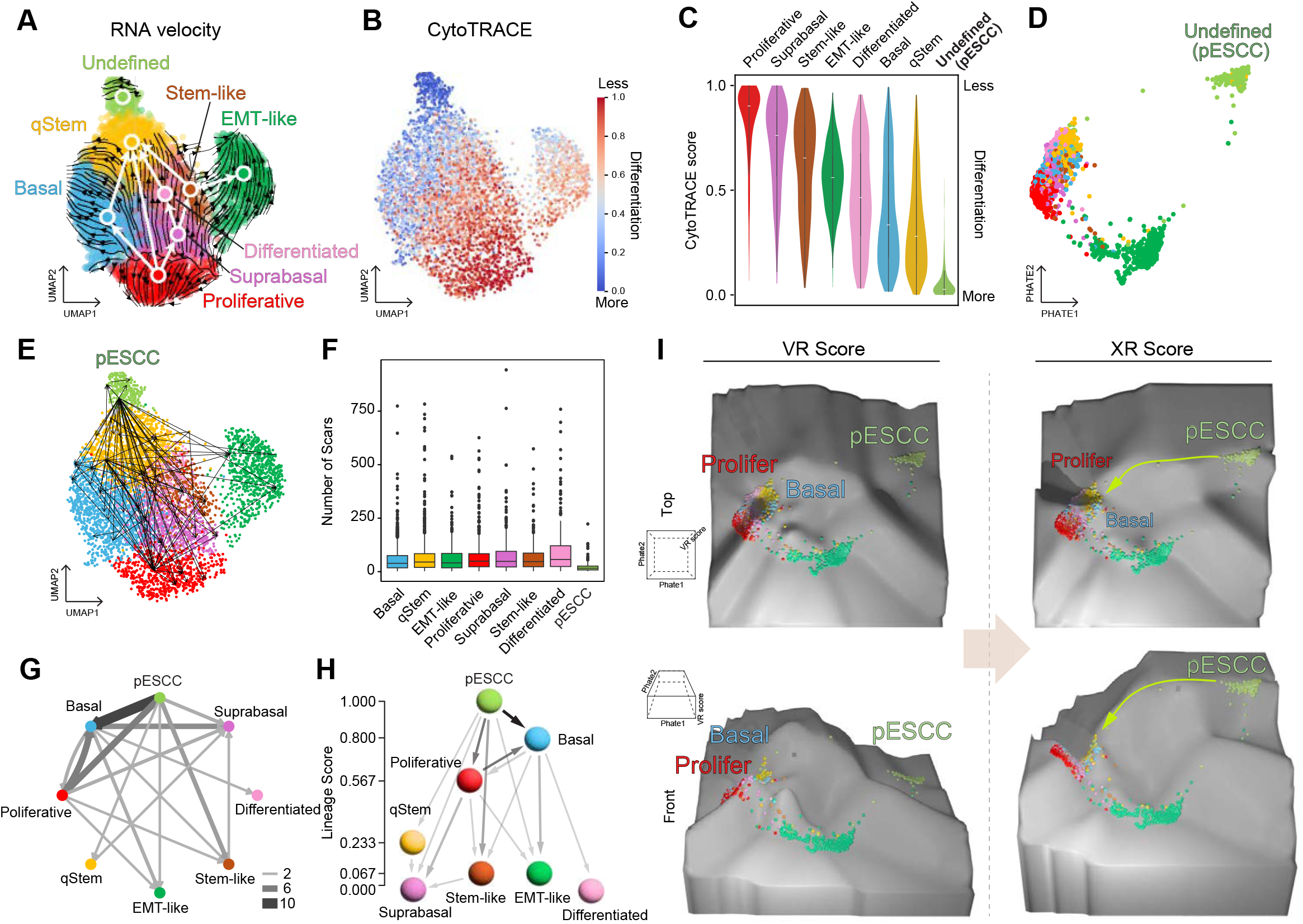
Identification of precursor cells through barcode-based cell tracing. **(A)** UMAP with RNA velocity analysis results depicting inferred cellular trajectories based on transcriptional dynamics. Arrows indicate predicted lineage progression. **(B)** UMAP displaying CytoTRACE analysis results, in which higher scores indicate lower differentiation states across cell populations. **(C)** Violin plots showing CytoTRACE score distribution across cell clusters. **(D)** PHATE visualization of cellular differentiation states. **(E)** UMAP illustrating differentiation pathways based on barcode and scar-based lineage tracing, highlighting directional transitions between cells with shared barcode information across clusters. **(F)** Box plot illustrating the distribution of CRISPR/Cas9-induced scars across cell populations. The box represents the interquartile range (IQR), with the median shown as a horizontal line. Error bars indicate [SD/SEM] to show data variability. **(G)** Network-based lineage analysis illustrating cellular differentiation pathways based on barcode and scar information, with edge width reflecting the number of differentiation events. **(H)** Schematic representation of differentiation flow, in which distances between cell types correspond to network scores. **(I)** Waddington landscape illustrating cell lineage dynamics, comparing VR and XR scores to depict differentiation trajectories.

### Construction of cellular lineage network based on barcode and scar information

To investigate cellular lineage, we used CytoTRACE and RNA velocity, scRNA-seq-based cell lineage trajectory inference tools (21, 38, 39). By using these inference-based cell trajectory analyses, we found that the cellular lineage originated from the proliferating cell cluster and transitioned toward other cell types, including basal, suprabasal, and differentiated cells (Fig. 2*A-C*). Notably, a strong lineage trajectory toward the EMT-like cell cluster was observed (Fig. 2*A*), a feature absent in normal esophageal epithelium. Although these analyses revealed somewhat precise cell trajectories, the undefined cluster exhibited an isolated and distinct pattern compared with the other clusters, suggesting a unique cellular trajectory that was not evident in the inference-based analyses (Fig. 2*A-C* and *SI Appendix*, Fig. S2*A*). A PHATE (potential of heat-diffusion for affinity-based trajectory embedding) map (40) further supported the distinct characteristics of the undefined cluster, as it remained separated from other clusters (Fig. 2*D*).

To identify the connections between the undefined cluster and other cells, we analyzed barcode and scar data and found 497 cellular lineage links between cells sharing the same barcode (Fig. 2*E*). Strikingly, the undefined cell cluster exhibited the lowest number of scar types, yet a substantial portion of lineage links from the undefined cluster was directed toward other cell types, with the strongest signals observed in the proliferating and basal cell populations (Fig. 2*E* and *F* and *SI Appendix*, Fig. S2*B* and *C*). This observation suggests that the undefined cells represent preneoplastic cells of esophageal squamous cell carcinoma (pESCCs) that give rise to other cell types, including proliferating and basal cells.

To further delineate the cellular lineage in preneoplastic cells, we constructed a cellular lineage network with barcode and scar-based links and calculated a network score based on the number of connections between clusters. This analysis revealed a strong trajectory from pESCCs to basal and proliferating cell clusters, with relatively weaker connections to other cell types (Fig. 2*G* and *H*). These results indicate that pESCCs are more closely related to proliferating and basal cells than to other cell populations.

Although the network score provided direct evidence of the pESCC lineage leading to proliferating and basal cells, the overall map of cell lineages and their sequential orders remained unclear. To address this, we applied the network score to the traditional valley-ridge (VR) score method (41, 42) and developed the eXamined Ridge (XR) score, as outlined in the Methods (*SI Appendix*, Fig. S2*D*). The XR score provided a more precise representation of the undefined cluster as progenitor cells with the highest cellular plasticity, which was visualized on Waddington’s landscape (Fig. 2*I* and *SI Appendix*, Fig. S2*E*). In contrast, the VR score, which relies on inference-based approaches, failed to capture the high plasticity of pESCCs. Moreover, our XR score more effectively distinguished other parts of the preneoplastic cell lineage, such as the basal/proliferating-suprabasal-EMT-like cell axis, than did the VR score in both PHATE (40) (*SI Appendix*, Fig. S2*E* and *F*) and uniform manifold approximation and projection (UMAP) embeddings (*SI Appendix*, Fig. S2*G* and *H*).

### Identification of pESCCs-specific markers

To identify molecular markers for detecting the progenitor cell cluster, we used differential gene expression analysis to compare the progenitor cell cluster with other clusters. Interestingly, the expression of marker genes in the progenitor cell cluster exhibited relatively low “fold-change” values compared with marker genes from other clusters, highlighting its quiescent, progenitor-like characteristics (Fig. 3*A*).

**Fig. 3.**
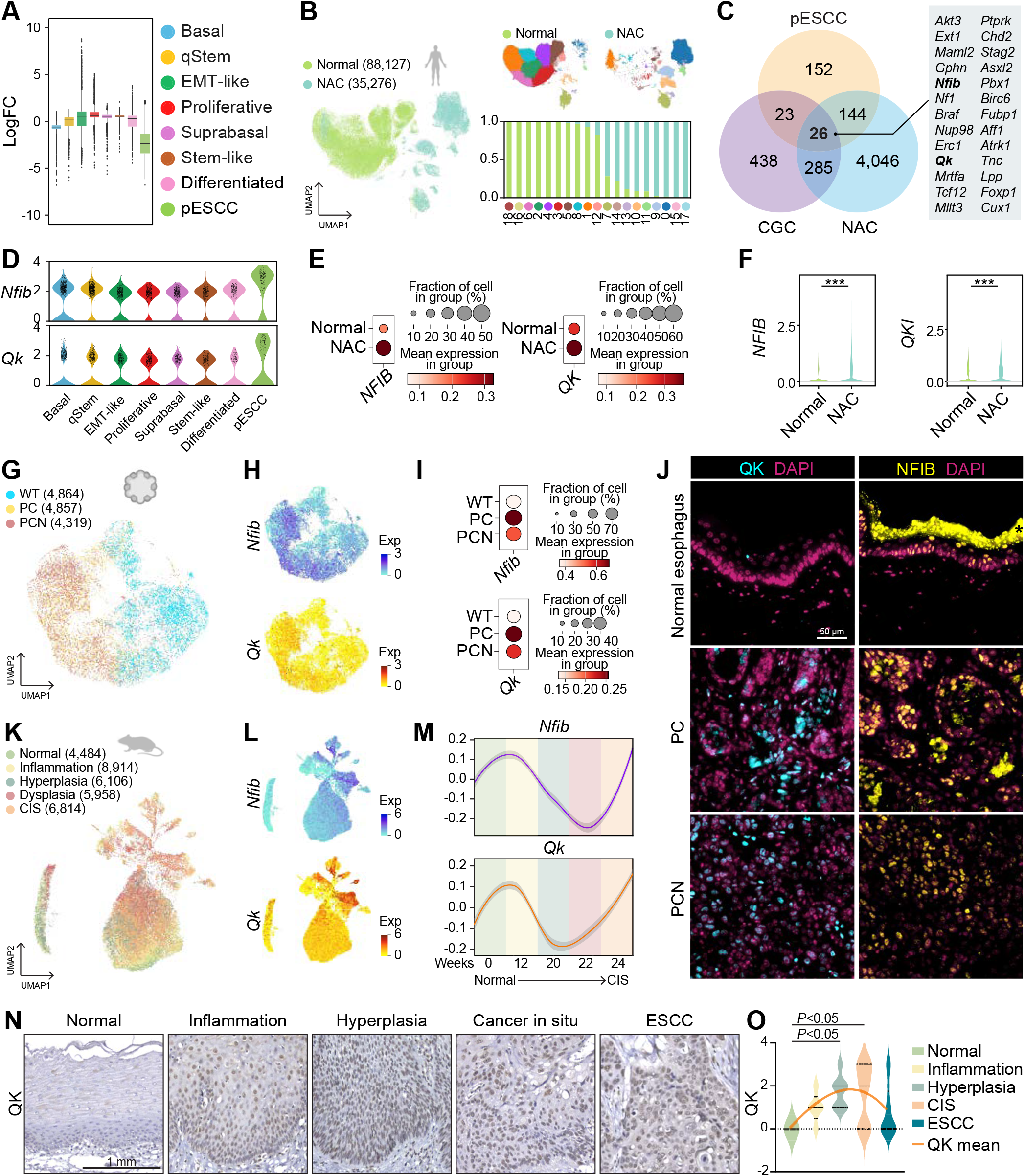
Expression of esophageal squamous cell carcinoma markers in mouse and human esophagus. **(A)** Box plot showing log (fold change) in significant markers (*P* < 0.05) across cell clusters. The box represents the interquartile range (IQR), with the median shown as a horizontal line. Error bars indicate [SD/SEM] to show data variability. **(B)** UMAP of healthy normal and normal adjacent cancer (NAC) human datasets. Proportions of normal and NAC cells in each cell cluster are displayed with bar plots. **(C)** Venn diagram illustrating the overlap among esophageal squamous cell carcinoma (pESCC) markers, NAC markers, and Cancer Gene Census (CGC) genes. A total of 26 genes are shared among all three groups. **(D)** Violin plots showing the expression profiles of *Nfib* and *Qk* across different cell clusters. **(E)** Dot plots of *NFIB* and *QKI* expression in human normal and NAC datasets. **(F)** Violin plots illustrating expression levels of *NFIB* and *QKI* across healthy normal and NAC human datasets. (Student’s *t*-test, \*\*\**P*□<□0.001). **(G)** UMAP of genetically engineered mouse esophageal organoid datasets. WT, wild-type; PC, *Trp53 Cdkn2a* double knockout (KO); PCN, *Trp53 Cdkn2a Notch1* triple KO. (**H-I**) Feature plots (H) and dot plots (I) of *Nfib* and Qk expression. **(J)** Immunostaining of *QKI* and *NFIB* expression in mouse esophagus tissue and allograft tumors from PC and PCN cells. Representative images are shown. Scale bars, 50 μm. **(K)** UMAP of carcinogen (4-NQO)-induced mouse esophageal tissue datasets. **(L)** Feature plots of *Nfib* and *Qk* expression in mouse tissue datasets. **(M)** Linear regression plot showing the expression levels of *Nfib* and *Qk* along the differentiation trajectory in mouse tissue datasets. (**N-O**) Representative IHC images (N) and quantification (O) of *QKI* expression in different stages of human esophageal disease. Scale bars, 1 mm (N). (Student’s *t*-test, \**P*□<□0.05).

To refine the identification of specific markers, we conducted additional scRNA-seq analysis on normal human esophageal epithelial cells and normal adjacent cancer (NAC) cells from patients with esophageal squamous cell carcinoma (ESCC) (43-46). NAC cells exhibit intermediate characteristics between normal and tumor cells and were therefore considered to include preneoplastic cells. As expected, NAC cells displayed several distinct cell clusters on a UMAP after integration with normal esophageal epithelial cells, suggesting that their cell identities had shifted from a normal state (Fig. 3*B*).

From the intersection of these analyses, we identified a set of potential markers. Among these, we specifically focused on genes with known oncogenic functions, as defined by the Cancer Gene Census (47), a comprehensive database cataloging genes with strong experimental and clinical evidence of involvement in cancer. The Cancer Gene Census classifies genes based on their roles as oncogenes or tumor suppressors, considering factors such as somatic mutations, germline predisposition, and functional studies. To refine our selection, we prioritized genes with established oncogenic activity, particularly those recurrently mutated, amplified, or overexpressed in cancer, leading to the identification of 26 candidate markers of pESCCs (Fig. 3*C* and *SI Appendix*, Fig. S*3*). Among these, *Nfib* and *Qk* were selected as consistent markers in both mouse and human models, not only because of their significantly elevated expression in pESCCs and NAC cells but also because high-specificity antibodies were available for their detection. This enabled precise experimental validation, which allowed accurate localization and functional characterization in preneoplastic cells (Fig. 3*C-F* and *SI Appendix*, Fig. S4*A* and *B*).

We further validated the increased expression profiles of *Nfib* and *Qk* by analyzing scRNA-seq data from various experimental models. First, we examined scRNA-seq data from 3D organoids of wild-type (WT) cells, *Trp53* and *Cdkn2a* double-KO (PC) cells, and *Trp53, Cdkn2a*, and *Notch1* triple-KO (PCN) cells. *Nfib* and *Qk* expression were upregulated in PC organoids but slightly decreased in PCN organoids (Fig. 3*G-I* and *SI Appendix*, Fig. S4*C*). These findings were further validated with immunohistochemical staining of normal esophageal tissue, PC-derived preneoplastic (benign) tumors, and PCN-derived neoplastic (malignant) tumors. Both QKI and NFIB expression were elevated in benign tumors but slightly reduced in neoplastic tumors (Fig. 3*J*). Because PC and PCN organoids represent preneoplasia and early neoplasia, respectively (27), these results suggest that *Nfib* and *Qk* are specifically upregulated during the preneoplastic stage of cancer progression.

We also analyzed scRNA-seq data from a carcinogen-induced mouse ESCC model. The expression of *Nfib* and *Qk* was upregulated during the early stages of cancer progression, especially in the inflammation stage. However, their expression gradually declined as the cells progressed toward neoplastic transformation and increased again at the cancer in situ stage (Fig. 3*K-M* and *SI Appendix*, Fig. S4*D*). This dynamic pattern was also consistent in human samples across different disease stages. A significant upregulation of NFIB and QKI expression was observed during the inflammation, hyperplasia, and cancer in situ stages, whereas a decline was noted in the later stages of ESCC (Fig. 3*N-O* and *SI Appendix*, Fig. S4*E* and *F*).

To expand on these findings, we performed unsupervised clustering to derive a gene expression signature for pESCCs. This analysis identified a signature, NMF7, comprising 531 genes, including 22 of the 26 previously defined pESCC markers (Fig. 3*C* and *SI Appendix*, Fig. S5*A* and Table S1). The NMF7 signature was significantly upregulated in human NAC tissues, suggesting the presence of a pESCC-like program in these samples (*SI Appendix*, Fig. S5*B-C*). In a mouse tumorigenesis model, its expression patterns mirrored those of Nfib and Qk, supporting NMF7 as a robust and transferable marker of preneoplastic cells across datasets (*SI Appendix*, Fig. S5*D-E*).

### pESCCs expansion during the early stage of tumorigenesis

Since our findings were from an in vitro model without any selection pressure, we investigated the dynamics of pESCCs within an intact microenvironment, where the immune niche is less tumor-favorable. We first performed spatial transcriptomics (Xenium Prime In Situ) to map the structural and spatial information of both PC-tumor and normal esophagus (Fig. 4*A*). After cell segmentation and annotation, we found that PC-derived tumors exhibited a well-differentiated architecture, composed of numerous units with centrally located keratin pearls surrounded by epithelial cells (Fig. 4*B*). Each unit displayed a cortex-to-core differentiation pattern, with basal cells at the periphery and differentiated cells at the core. This organization suggests that basal cells primarily interact with immune cells, and thus, resistance to immune surveillance may be critical for their persistence. We then compared the expression of the pESCC marker, *Nfib*, in normal and PC-tumor tissues. *Nfib* expression was significantly elevated in the PC-tumor (Fig. 4*C*) and was predominantly localized to the basal cells (Fig. 4*D*). In addition, a subset of *Nfib*+ cells in the PC-tumor showed co-expression of *Mki67*, indicating a proliferative phenotype. This basal cell-focused and proliferative phenotype was confirmed in the normalized expression values (Fig. 4*E* and *F*). *Qki* was not assessed due to the lack of its probes in the Xenium 5K geneset. These data suggest that pESCCs constitute a dominant and actively proliferating cell population at the tumor-immune interface.

**Fig. 4.**
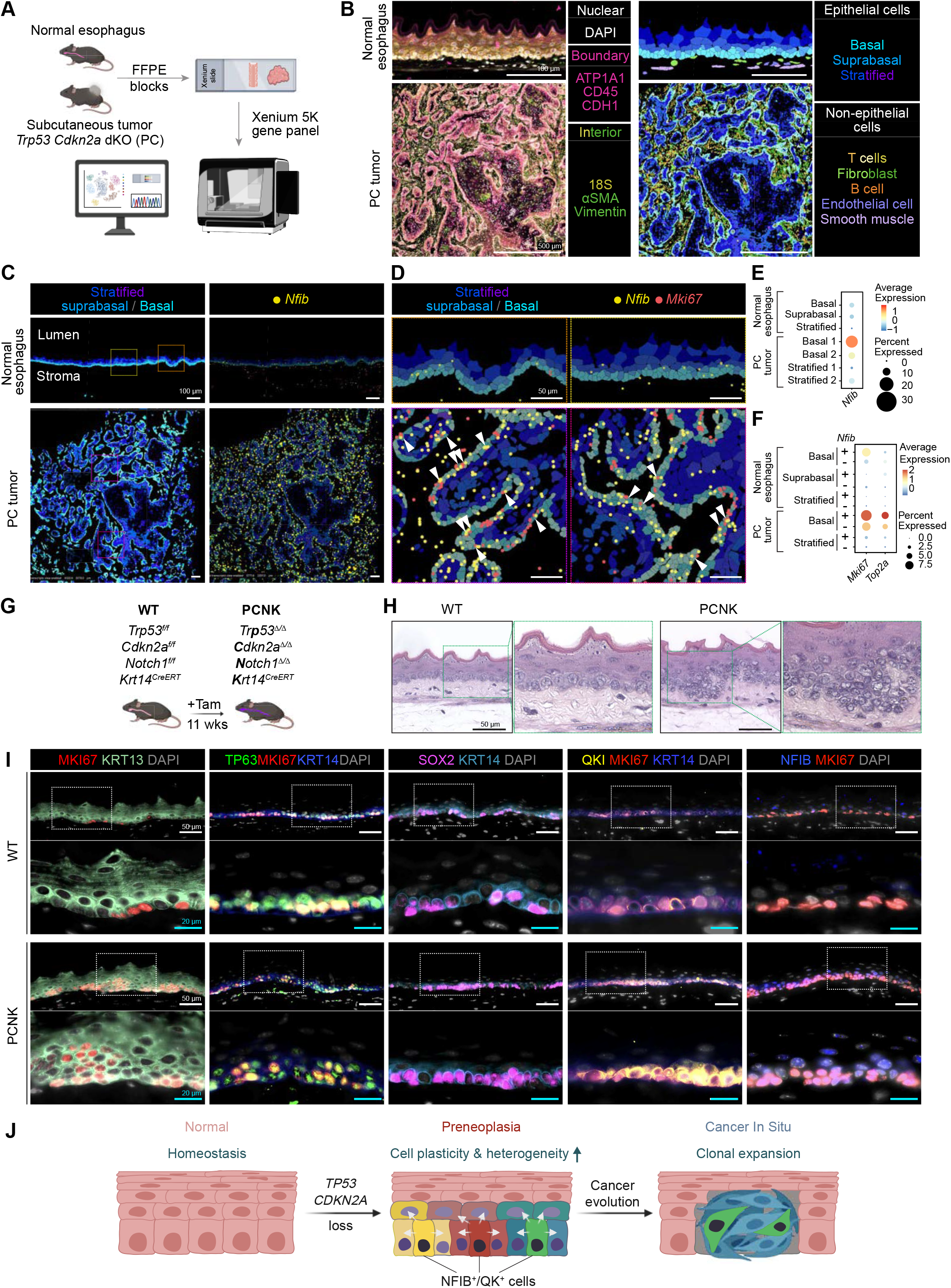
pESCC expansion during early tumorigenesis. (**A**) Workflow of spatial transcriptomics (Xenium In Situ) analysis using normal esophagus and PC-derived tumor. (**B-D**) Cell segmentation and annotation of Xenium In Situ data (B). (C-D) Epithelial cells showing *Nifib* expression at low magnification (C) and *Nfib* and *Mki67* expression in regions of interest (ROIs) (D); Arrowheads indicate cells co-expressing both genes. Scale bars, 500 μm (B, bottom panels), 100 μm (B, upper panels, C) and 50 μm (D). (**E**) Normalized *Nfib* expression in each epithelial cell type. (**F**) Normalized *Mki67* and *Top2a* expression in Nfib+ versus Nfib- epithelial cell groups. (**G**) Schematic of the PCNK mouse model. (**H**) H&E staining of WT and PCNK mouse esophagi. Representative images are shown. Scale bars, 50 μm (**I**) Immunofluorescent staining of WT vs. PCNK esophageal samples for markers of proliferative (MKI67), suprabasal (KRT13), basal (TP63, SOX2, and KRT14), and pESCC (QKI and NFIB) cells. Representative images are shown. Scale bars, 50 μm (low magnification) and 20 μm (high magnification). (**J**) Schematic representation of esophageal squamous cell carcinoma expansion during cancer progression.

To evaluate pESCC behavior in the esophageal immune context distinct from subcutaneous tumors, we additionally used a genetically engineered mouse model (GEMM) to induce preneoplastic lesions. We established a compound mouse strain for triple-KO of *Trp53, Cdkn2a*, and *Notch1* using the *Krt14*^CreERT^ driver. Via tamoxifen administration, the *Trp53, Cdkn2a*, and *Notch1* alleles were ablated in *Krt14*- expressing epithelial cells (Fig. 4*G*). 11 weeks later, mice were euthanized for esophageal tissue analysis. Histological analyses of the PCNK (*Trp53*^*Δ/Δ*^ *Cdkn2a*^*Δ/Δ*^ *Notch1*^*Δ/Δ*^ *Krt14*^*Cre/ERT*^) esophagus displayed hyperplastic phenotypes, including increased epithelial cell numbers and hyperchromatic nuclei (Fig. 4*H*). This hyperplastic phenotype was confirmed by the expression of proliferation (MKI67) and basal stem cell (TP63, SOX2) markers (Fig. 4*I*). However, these proliferative or basal cell markers were not specific to the hyperplastic lesions since they reflect the number of basal cells in both normal and PCNK esophagi. In contrast, pESCC markers identified in our study, QKI and NFIB, were significantly increased in the hyperplastic lesion while nearly absent in the normal esophagus.

Together, these findings indicate that pESCCs survive and expand during preneoplastic or early neoplastic stages of ESCC despite immune-mediated selection pressure. Notably, while pESCCs occupy only a small fraction of cells in scRNA-seq datasets, they represent a substantially larger proportion under in vivo selection pressures. However, as a unit in the tumor mass enlarges, pESCCs shift toward the periphery, while the immune cell-cleared core becomes occupied by other epithelial cell types. These findings also imply that pESCCs might be essential for overcoming selection pressures during the initial stages of tumor development but become less critical for clonal expansion (Fig. 4*J*).

## Discussion

This study advances our understanding of esophageal preneoplasia by identifying preneoplastic progenitor cells (pESCCs) and their cellular dynamics. By combining genetic barcoding with scRNA-seq, we established a comprehensive cellular lineage map of esophageal preneoplastic cells. This approach allowed us to overcome key limitations of scRNA-seq-based lineage analysis, which primarily relies on inference-based predictions or prior biological knowledge. Although inference-based analyses have been effective in delineating cell lineage trajectories in normal tissues, they often overlook unknown cell types that are critical to transformed cell populations owing to the lack of direct experimental validation. Here, we identified lineage connections through experimental validation, revealing a specific cell cluster functioning as a progenitor population that gives rise to proliferating and basal cells. This finding was further reinforced by the development of the eXamined Ridge (XR) score, which highlighted the distinct role of this progenitor cluster in driving cellular heterogeneity during early neoplastic transformation.

Preneoplastic cells have gained increasing attention because of the transcriptional and epigenetic reprogramming that occurs at this stage, even though these cells remain phenotypically normal (48, 49). During this early phase of tumorigenesis, cells acquire plasticity, leading to the emergence of distinct progenitor cell populations (8). Most of the recent studies on precancerous stages of esophageal squamous cell carcinoma (ESCC) have focused on dysplastic stages, later points in disease progression, making them less suitable for understanding the initial events of tumorigenesis (50-52). In contrast, our study focused on earlier cellular events, where only two genetic alterations (*Trp53* and *Cdkn2a* loss) were introduced. Although these genetic alterations are commonly observed in patients with ESCC, research on them has been limited, as they alone are insufficient to drive neoplastic transformation. However, as shown in Figure 3, *Trp53* and *Cdkn2a* loss (the PC model) was sufficient to generate pESCCs, which exhibited the highest cellular plasticity.

Intriguingly, pESCCs displayed unique features, including suppressed mRNA splicing and high activation of Mtf1, Zbtb24, Tlx3, and Thap1 regulons. Because these regulons have critical roles in tumor progression, self-renewal, and DNA methylation, it is highly plausible that pESCC clusters serve as a key source of cellular plasticity in early ESCC tumorigenesis. Functional analysis of the downstream targets of these TFs supports a model of transcriptional poising. Although the cells exhibit functional arrest associated with abnormalities in mRNA splicing, these TFs are linked to large-scale transcriptional reprogramming and activation of pathways preparing the cells for rapid future responses (34-37). We also identified two transcriptional markers, *Nfib* and *Qk*, that were enriched in pESCC clusters and had critical roles in cancer progression. These markers were validated across several experimental models, including 3D organoids, mouse tissue models, and human ESCC samples. Interestingly, their expression patterns were dynamic, showing upregulation during early preneoplasia, downregulation during cellular expansion, and re-emergence during cancer in situ (Fig. 3*M-O*). Eventually, their expression slightly decreased again in later ESCC stages (Fig. 3*M-O*). Although the dynamics of pESCCs are not entirely consistent between mouse and human temporal samples, a clear trend emerged: pESCCs are significantly expanded during the early stages of both preneoplasia and neoplasia. This expansion suggests that pESCCs provide the cellular plasticity required to overcome early selection pressures, ensuring their survival and adaptation during the initial stages of tumorigenesis.

The dynamic expression pattern of *NFIB* and *QKI* suggests that they may have dual roles in early preneoplasia and neoplasia. Given the known functions of NFIB as a transcription factor and QKI as an mRNA splicing regulator, their roles are likely crucial for modulating gene expression that is required for cellular adaptation to the microenvironment during these early stages. Further studies are needed to elucidate their precise regulatory mechanisms across different stages of preneoplasia and cancer progression.

Beyond pESCCs, we also identified an EMT-like cell cluster, which was absent in the normal esophagus (Fig. 2). Interestingly, this EMT cluster originated from progenitor cells (proliferating or basal cells) or pESCCs but not from differentiated cells (Fig. 2*G*). This suggests that the emergence of mesenchymal-like differentiation in preneoplastic esophageal cells was previously unrecognized. Our findings indicate that preneoplastic esophageal cells diversify cell lineages, generating cellular heterogeneity beyond the normal epithelial trajectory.

Despite the homogeneous genetic alterations in our preneoplasia model (*Trp53* and *Cdkn2a* loss), we observed highly variable and heterogeneous cell fates (*SI Appendix*, Fig. S4*G* and *H*). This observation suggests that cancer evolution in preneoplastic cells is not solely governed by genetic changes but is also influenced by non-genetic mechanisms, such as epigenetic factors and tumor niche (53, 54). Additionally, the tumor microenvironment, including immune cells, may impose selection pressures on tumors (55, 56). To address this, we generated a PCNK mouse model that also showed the specific expression of NFIB and QKI in the hyperplastic lesions (Fig. 4). Similarly, the histological classification of pESCCs remains undefined. However, animal studies by Chang et al. and Yao et al. provide relevant insights (57, 58). In our model, pESCC marker expression peaks at 12 weeks post-4-NQO treatment, preceding the 16-week low-grade intraepithelial neoplasia (LGIN) stage reported in these studies (57, 58). This aligns with human IHC data showing higher expression during inflammation and hyperplasia than in neoplastic stages. These findings suggest that pESCCs emerge in early, histologically normal or preneoplastic stages not fully captured by current classifications. While our study provides a detailed snapshot of preneoplastic cells, their roles in cancer evolution remain to be fully explored. Future research might include assessing the tumorigenicity of pESCC by sorting these cells for syngeneic transplantation or performing cell ablation of pESCC in the PCNK model.

In summary, our study provides a comprehensive cellular and molecular framework for understanding esophageal preneoplasia by identifying a progenitor cell cluster with distinct transcriptional signatures. Our findings highlight the dynamic cell lineage relationships that drive early cancer progression, offering a foundation for future investigations into tumor initiation. By uncovering key molecular markers associated with preneoplastic transformation, this study also provides valuable insights for refining ESCC risk stratification and developing early diagnostic strategies.

## Ethics Approval and Consent to Participate

All animal experiments were performed under an Institutional Animal Care and Use Committee (IACUC)-approved protocol, adhering to the guidelines established by the Association for the Assessment and Accreditation of Laboratory Animal Care (AAALAC). To ensure compliance with ethical standards and species-specific care requirements, animals were procured and housed exclusively through the Division of Veterinary Medicine and Surgery (DVMS).

## Consent for Publication

Not applicable.

## Competing interests

The authors declare no competing interests.

## Funding

This work was supported by the National Cancer Institute (CA286761 to K.-P.K.; CA193297, CA278971, CA279867, CA256207, and CA278967 to J.-I.P.). The core facilities at MD Anderson received support from the National Cancer Institute Cancer Center Support Grant (P30 CA016672). The core facilities at Baylor College of Medicine were funded by the Cancer Prevention and Research Institute of Texas (RP180672, RP200504) and the National Institutes of Health (CA125123, RR024574).

## Authors’ Contributions

J.J.: Conceptualization, methodology, investigation, software, analysis, data curation, writing (original draft), visualization; K.-P.K.: Conceptualization, methodology, investigation, software, analysis, data curation, writing (original draft), visualization, funding acquisition; J.Z.: Investigation; S.J.: Investigation; J.-I.P.: Conceptualization, methodology, analysis, writing (review and editing), visualization, supervision, project administration, funding acquisition

## Acknowledgements

We thank Christine F. Wogan (Division of Radiation Oncology, MD Anderson) for manuscript editing. The core facilities at MD Anderson (DNA Sequencing and Genetically Engineered Mouse Facility) were instrumental in supporting this work, with additional assistance from the Cytometry & Cell Sorting Core and Single Cell Genomics Core at Baylor College of Medicine. Graphical illustrations were created using BioRender.com.

## Data and code availability

scRNA-seq and Xenium In Situ data are available via the GEO database (GSE289049; log-in token for reviewers: azgbasusfhkxhah and GSE304762; log-in token for reviewers: ytkvouqubtqpzst). The code used to reproduce the analyses described in this manuscript can be accessed via GitHub (https://github.com/jaeilparklab/Esophagus_PNPC) and will be available upon request.

## Materials and Methods

### Cell culture

PC (*Trp53 Cdkn2a* double knockout [KO]) and PCN (*Trp53 Cdkn2a Notch1* triple KO) organoids and 2D cells were prepared as previously described (27). Briefly, esophageal epithelial cells isolated from *Trp53*^floxed/floxed^ mice were used to establish esophageal organoids. Ad-Cre-EGFP (University of Iowa) was introduced to develop Trp53^−/−^ organoids. Next, CRISPR/Cas9 system-mediated *Cdkn2a* or *Notch1* KO was added to the Trp53^−/−^ organoids to establish the PC and PCN organoids. Organoid cells were dissociated into single cells and cultured on 24-well plates with Dulbecco’s Modified Eagle Medium (DMEM) + 10% fetal bovine serum (FBS) with 10 μM Y-27632. After the third passage, cells were cultured without Y-27632.

### Barcode library preparation

Barcode libraries were prepared as described elsewhere (29). iTracer plasmid was purchased from the European Plasmid Repository. Barcode sequences were amplified by using a custom-designed barcode template and primers: Barcode-template_perturb, 5’- ACGAGCTGTACAATAAGCTGATCCWNNNNNNNNNWCCAAGCGCCCTTGAGCATCTG-3’; Barcode forward, 5’-GGACGAGCTGTACAAGTAAGCTG-3’; Barcode reverse, 5’- GAATCAGATGCTCAAGGGCGCTTG-3’. Amplified barcodes were purified and ligated with NotI-linearized iTracer plasmid by using an NEBuilder HiFi DNA Assembly kit. The ligation product was transformed into DH5α cells, which were spread on LB plates containing ampicillin and incubated overnight at 37°C. Colonies were then scraped from the LB plates and pooled in a 2× YT medium for plasmid extraction by using the EndoFree Plasmid Maxi Kit (Qiagen).

### Transfection and sequencing library preparation for scRNA-seq

PC cells with constitutive expression of Cas9 were transfected with a prepared barcode library by using Lipofectamine 3000. After transfection, cells were amplified for three passages over six days and subjected to flow cytometry and scRNA-seq. For flow cytometry, 1 million RFP-positive and live cells were sorted and prepared for scRNA-seq. Targeted cell recovery was approximately 20,000 cells. After the cDNA library was prepared, the gene expression library was made by using a 10x Genomics 3’ v2 kit (Baylor College of Medicine), and barcode and scar libraries were generated separately. The barcode region was amplified by 27 cycles of root PCR (Forward, 5’-GACGACGGCAACTACAAGACC- 3’; Reverse, 5’-CTACACGACGCTCTTCCGATCT-3’) and 10 cycles of nested PCR for Barcode (Forward, 5’-GTGACTGGAGTTCAGACGTGTGCTCTTCCGATCTGGATCACTCTCGGCATGGA-3’; Reverse, 5’- CTACACGACGCTCTTCCGATCT-3’) and Scar (Forward, 5’- GTGACTGGAGTTCAGACGTGTGCTCTTCCGATCTGGCCAACTTCAAGATCCGCC-3’; Reverse, 5’- CTACACGACGCTCTTCCGATCT-3’) regions. The PCR product was purified by using 1× AMPure XP beads (Beckman). Indexing PCR for barcode and scar was conducted for 14 cycles by using dual-index primers from the 10x Genomics. Indexed libraries were purified by using 0.6× SPRI [solid-phase reversible immobilization] bead purification and quantified. The three libraries were sequenced for gene expression (400M reads), scar regions (167M reads), and barcode sequences (100M reads) on an Illumina NovaSeq (Novogene), and the fastq files were mapped to the customized reference genome by using CellRanger (10x Genomics Cloud).

### Preprocessing scRNA-seq data

scRNA-seq data were preprocessed by using Scanpy (version 1.10.4) (59). Quality control (QC) steps were applied to filter cells and genes with low expression levels. Specifically, cells expressing fewer than 100 genes were removed, and genes detected in fewer than three cells were filtered out. After QC, only cells with extractable barcode information from the alignment BAM files were selected for downstream analysis. This ensured that the dataset included cells with both transcriptomic and lineage information, which is critical for accurate lineage reconstruction. For subsequent analyses, including the identification of new cell populations and their unique markers, all genes passing the QC criteria were used to ensure a comprehensive and unbiased discovery process.

### Regulon Analysis and Visualization

Regulon analysis was performed using pySCENIC (60) to infer transcription factor-target interactions and assess regulon activity at the single-cell level. Network inference was performed using GRNBoost2, followed by motif enrichment analysis with RcisTarget to refine the inferred networks, and AUCell to quantify regulon activity. The resulting regulon activity matrix was imported into R for visualization, where the reshape2 package (61) was used to transform data into a suitable format, ComplexHeatmap (62) was used to generate clustered heatmaps, and ggplot2 facilitated violin plots, boxplots, and correlation analyses. Statistical significance was assessed using t-tests, enabling identification of key transcriptional regulators driving cellular heterogeneity.

### Gene Set Enrichment Analysis

Gene set enrichment analysis was performed using the fgsea package (63) in R to identify significantly enriched gene sets. Differential expression analysis results were ranked using the score provided by scanpy’s rank_genes_groups function, and the pre-ranked gene list was used as input for fgsea. The msigdbr package was employed to retrieve molecular signature gene sets, including hallmark and curated pathways, while biomaRt (64) was used for gene annotation and conversion between different gene identifiers to ensure compatibility with the selected gene sets.

### Barcode and scar analysis with LINNAEUS

Barcode sequences were analyzed with the LINNAEUS system by amplifying the regions flanking the barcode, including upstream and downstream sequences, and aligning them to the reference genome (mm10) by using the web-based Cell Ranger platform (65). Cells were assigned to the same lineage only if they possessed identical barcode sequences, ensuring high specificity in lineage reconstruction. For each pair of cells with the same barcode, scar information was extracted from the alignment BAM files by using Samtools (version 1.21) (66), which analyzed insertion and deletion patterns encoded in the CIGAR string. The accumulation of scars was compared between cell pairs to infer differentiation directionality, with the assumption that scars progressively accumulate in more differentiated cells. This approach provided detailed insights into lineage relationships and cellular trajectory within the population.

### Cell lineage inference

To infer cellular lineage trajectories, several complementary approaches were used. Cell cycle phases were inferred by using a curated list of cell cycle markers as described elsewhere (67), enabling the identification of cells in G1, S, or G2/M phases. Pseudo-time analysis was conducted by using CytoTRACE(21) to estimate differentiation potential along the inferred trajectory (21). RNA velocity analysis was performed with Velocyto (38) and scVelo (39), which provided dynamic insights into cell state transitions based on spliced and unspliced mRNA ratios. To further analyze the directionality and entropy of cellular transitions, SCENT(22) was used to calculate cellular entropy, which was integrated with RNA velocity data to compute a VR score (41). This VR score quantified differentiation potential and lineage progression for each cell cluster. Finally, a Waddington-like landscape was visualized by using Houdini, where the VR score was plotted along the vertical axis to represent differentiation potential, and the pseudo-time coordinates were mapped onto PHATE (40) coordinates representing pseudo-temporal progression on the horizontal axes with the VR score on the vertical axis. This integrated approach provided a detailed view of lineage relationships and the differentiation dynamics within the cell population.

### Identification of molecular markers of pESCCs

To identify molecular markers specific to pESCCs, we conducted integrative analyses of scRNA-seq datasets from mouse cell lines and human esophageal cancer patient samples. In the mouse dataset, genes upregulated in the pESCC cluster were compared with other clusters to generate a PPC-specific gene list. For human data, samples from patients with esophageal cancer were subjected to scRNA-seq analysis. Differentially expressed genes were identified by comparing NAC samples with normal tissue. The PPC-specific gene list from the mouse dataset was intersected with the NAC-derived differential gene expression list to identify conserved candidate markers across species. To prioritize markers with oncogenic relevance, the resulting list was cross-referenced with the oncogenic gene list defined by the Cancer Gene Census (47). This analysis identified 26 candidate pESCC markers, representing conserved molecular signatures with potential roles in early esophageal cancer progression. These markers were subsequently used for further validation and functional studies.

### Calculation of XR scores

XR scores were calculated by integrating the VR score and the Network Score (NS) to represent lineage dynamics and differentiation potential. The NS was derived from a lineage network constructed from barcode and scar information. For each cluster, the NS was defined as the number of differentiation cell pairs directed toward other clusters within the network, quantifying the cluster’s contribution to lineage transitions.

Both the VR score and NS were normalized to ensure comparability across clusters by using the following formulas:

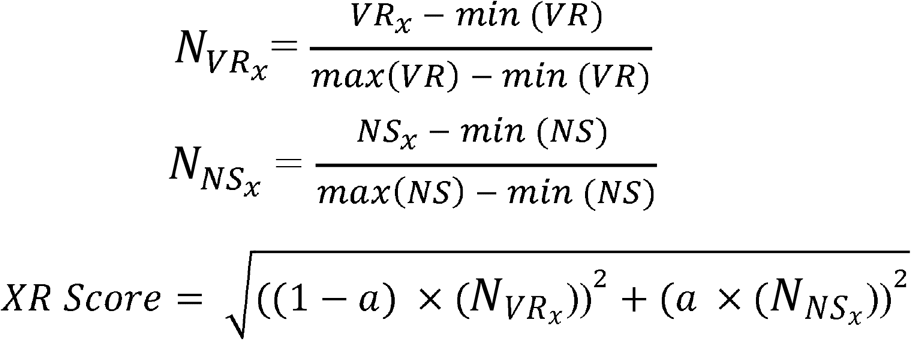

The XR score was then calculated as:

Here, *a* is a weighting coefficient that balances the contributions of the VR score and NS. To determine the optimal *a*, coefficients were tested from 0.5 to 0.9 in increments of 0.1. For each coefficient, the standard deviation of XR scores across clusters was calculated. Then, the median of these standard deviations across all clusters was computed to produce a single value for each coefficient. The coefficient that resulted in a median standard deviation of at least 0.01 was selected, ensuring consistent and robust differentiation patterns while avoiding excessive homogeneity in XR scores (*SI Appendix*, Fig. S2*D*). The resulting XR scores were visualized by using Houdini to create a Waddington-like landscape, with the XR score plotted on the vertical axis and PHATE(40) coordinates representing pseudo-temporal progression on the horizontal axes. This visualization enabled a detailed interpretation of cellular differentiation trajectories and lineage relationships.

### Human and mouse scRNA-seq dataset analyses

Mouse esophageal tissue datasets were obtained from the authors of the previous study (58). Human healthy normal esophageal tissues and NAC datasets were downloaded from the Human Cell Atlas Data Portal (PRJEB31843) and National Center for Biotechnology Information Sequence Read Archive public databases (PRJNA828562, PRJNA672851, and PRJNA777911) (43-46).

### Allograft transplantation and tissue isolation

Five-week-old C57BL/6 mice were maintained in the Department of Veterinary Surgery and Medicine at MD Anderson. 2D-cultured PC or PCN cells (3 × 10^6^) were injected subcutaneously into the right dorsal flanks of mice. Animals were euthanized, and PCN and PC tumors were collected 100 days and 140 days after transplantation, respectively. The excised tumors were fixed and paraffin-embedded for immunostaining. To isolate normal esophagus tissue, 8-week-old mice were euthanized and subjected to cervical dislocation. Longitudinally opened esophagi were washed with phosphate-buffered saline (PBS), fixed, and paraffin-embedded. All animal procedures followed the guidelines of the Association for the Assessment and Accreditation of Laboratory Animal Care and institutional (MD Anderson) approved protocols. This study was compliant with all relevant ethical regulations regarding animal research.

### Xenium In Situ

Formalin-fixed, paraffin-embedded (FFPE) sections from normal esophagus and PC-derived tumors were mounted on Xenium-compatible slides in accordance with the manufacturer’s instructions. The experiments were performed at the Baylor College of Medicine Single Cell Genomics Core Facility using the Xenium In Situ platform (10x Genomics). A Xenium Prime 5K Mouse Pan Tissue & Pathways Panel was employed to profile transcripts from epithelial, immune, and stromal compartments. After probe hybridization and rolling circle amplification, iterative fluorescent labeling was carried out to decode transcript identities and spatial positions. Imaging and base calling were performed with the Xenium Analyzer, and initial data processing was completed using the Xenium Onboard Analysis Software. Cell segmentation was guided by multimodal staining. Gene expression matrices and spatial maps were first explored in Xenium Explorer, followed by downstream analyses, such as cell type annotation and clustering, conducted in Seurat (R).

### Mice

To generate the PCNK mouse, the *Krt14*^CreERT^ (Tg[KRT14-cre/ERT]20Efu; Jackson Laboratory; RRID: IMSR_JAX:005107) Cre driver strain was bred with mice carrying the alleles *Trp53*^f/f^ (B6.129P2- *Trp53*^*tm1Brn*^*/J*; Jackson Laboratory; RRID: IMSR_JAX:008462), *Notch1*^*f*/f^ (*Notch1*^*tm2Rko*^/GridJ; RRID: IMSR_JAX:006951), and *Cdkn2a*^f/f^. *Notch1*^f/f^ and *Cdkn2a*^f/f^ mice were kindly provided by Dr. Ronald A. DePinho (68) and Dr. Yejing Ge, respectively.

### Tamoxifen administration and tissue isolation

To induce Cre recombination, tamoxifen (Sigma-Aldrich) was dissolved in corn oil at a concentration of 20□mg/mL and administered intraperitoneally to 6-week-old mice at a dose of 75□mg/kg daily for three consecutive days. Control mice received intraperitoneal injections of corn oil alone. Mice were sacrificed 11□weeks after tamoxifen treatment. Esophageal tissues were collected as previously described, fixed overnight in 4% formaldehyde, and paraffin-embedded.

### Immunohistochemical analysis

For human esophagus analysis, tissue microarray slides from patients with disease at different stages were purchased from TissueArray.com (Cat# ES809 and ES804). Immunostaining was done as previously described (27). Briefly, paraffin-embedded tissue antigens were retrieved with a basic antigen retrieval buffer and a heat-induced method with a pressure cooker. Tissues were then blocked with goat serum in PBS and incubated with primary antibodies (QKI [Proteintech, 13169-1-AP, 1:200], NFIB [Proteintech, 29898-1-AP, 1:200], KRT13 [abcam, ab92551, 1:250], MKI67 [Invitrogen, 14-5698- 82, 1:200], SOX2 [Cell Signaling Technology, #3579, 1:250], TRP63 [abcam, ab124762, 1:250], KRT14 [abcam, ab7800, 1:250]). The immunohistochemical staining results were scored by three independent evaluators, and the results were analyzed and visualized by using GraphPad Prism (v10).

### Non-negative Matrix Factorization (NMF) Analysis

To identify robust underlying gene expression programs, we performed an unsupervised clustering analysis on the expression profiles of the top 3,000 most variable genes across all cells. We applied the NMF algorithm for this purpose. To determine the optimal number of factors (k), we tested ranks ranging from k=3 to k=10. We selected k=7 as the optimal factorization, as this value minimized the standard deviation of the gene count within each factor, indicating the most stable solution. The resulting analysis identified seven distinct gene expression programs (NMF1-7). The program most specifically associated with the pESCC population, NMF7, was defined as a gene signature. This signature comprises the 531 genes that contributed most strongly to the NMF7 factor. The complete list of genes in this signature is provided in *SI Appendix*, Table S1.

## Notes

### Competing Interest Statement

The authors have declared no competing interest.

### Summary of Updates

New experimental results from spatial transcriptomics and a genetically engineered mouse model were added as Figure 4. Instead, the previous Figures 1 and 2 were merged into Figure 1. Accordingly, Supplementary Figure and information were also updated.

