## Supplementary information for "Deciphering Precursor Cell Dynamics in Esophageal Preneoplasia via Genetic Barcoding and Single-Cell Transcriptomics"

**This PDF file includes:**

Figures S1 to S5
Tables S1

**Fig. S1. Characterization of Preneoplastic Esophageal Cell Populations**

(**A**) Violin plots illustrating expression levels of key marker genes across different preneoplastic cell clusters.
(**B-C**) UMAP (left) and violin plot (right) displaying mRNA splicing pathway scores (**B**) and mRNA splicing minor pathway scores (**C**) for individual cells, highlighting variation across cell populations.
(**D**) Receiver Operating Characteristic (ROC) curves evaluating the performance of the four key TFs in discriminating the 'Undefined' cell cluster from other cell types. The curves for *Mtf1*, *Zbtb24*, *Tlx3*, and *Thap1* were generated using their respective regulon activity scores. The Area Under the Curve (AUC) value for each TF is provided in the legend.
(**E**) Bar plots showing the results of a Gene Ontology functional enrichment analysis for Biological Processes, performed using the DAVID database. The analysis was conducted on the downstream target genes for each of the four key TFs (*Mtf1*, *Zbtb24*, *Tlx3*, and *Thap1*). The plots display the most significantly enriched biological processes, with bar length corresponding to the statistical significance (-log(P-value)).

**Fig. S2. Validation of Lineage Tracing and XR Score Calculation**

(**A**) PHATE visualization showing cellular differentiation states inferred by CytoTRACE.
(**B-C**) PHATE (**B**) and UMAP (**C**) visualizations illustrating the number of distinct scar types per cell.
(**D**) Standard deviation analysis of XR Score weight coefficients (0.5–0.9). The optimal coefficient was determined based on minimizing variance across clusters while preserving differentiation patterns.
(**E-F**) PHATE projection comparing XR Score (**E**) and VR Score (**F**).
(**G-H**) UMAP projection comparing XR Score (**G**) and VR Score (**H**).

**Fig. S3. Expression Patterns of Key Marker Genes in pESCC**

(**A**) UMAP projection (left) and dot plots (right) illustrating the expression levels of top marker genes in pESCC, highlighting their distinct transcriptional profiles.

**Fig. S4. Validation of findings using independent datasets from patient samples, organoid models, and mouse models.**

(**A-B**) UMAP projection (left) and violin plots (right) of *NFIB* and *QKI* expression in normal human esophageal epithelial cells, as well NAC cells from ESCC patients.
(**C**) Violin plots illustrating expression of *Nfib* and *Qk* in WT, PC, and PCN organoid models.
(**D**) Violin plots displaying *Nfib* and *Qk* expression across different mouse model conditions, including normal, inflammation, hyperplasia, dysplasia, and CIS.
(**E**) Immunostaining of NFIB in normal, inflammatory, hyperplastic, and ESCC tissues. Scale bar: 1 mm.
(**F**) Violin plot quantifying NFIB IHC results from (**E**), comparing NFIB expression across different tissue states (Student's *t*-test, $*P<0.05$).
(**G**) UMAP projection visualizing CNV scores from CNV analysis, including PC cell clusters alongside normal and CIS mouse tissues.
(**F**) Heatmap illustrating CNV profiles across PC cell clusters, normal tissues, and CIS mouse tissues.

**Fig. S5. Identification of a pESCC-specific signature via NMF.**

(**A**) The process for determining the optimal number of NMF groups (k). The top-left panel shows the distribution of gene counts per group (boxplot) and the standard deviation (SD) of gene counts across groups for k ranging from 3 to 10. A value of k=7, which yielded the lowest SD, was selected for subsequent analysis (red vertical line). The remaining seven panels display the score of each NMF factor (NMF1-NMF7) as a feature plot on the UMAP representation and a corresponding violin plot across all cell clusters.

(**B**) Feature plot of the NMF7 score on human NAC and Normal scRNA-seq data, validating NMF7 as a pESCC-specific signature in human in vivo data.
(**C**) Violin plot of the NMF7 score, demonstrating a significant upregulation in NAC samples compared to Normal samples (***$P < 0.001$).
(**D**) Feature plot of the NMF7 score across a mouse in vivo tumorigenesis model, validating the NMF7 signature in the mouse model.
(**E**) Violin plot of the NMF7 score across different stages of cancer progression, revealing a dynamic pattern consistent with the expression of our previously identified pESCC markers, *Nfib* and *Qk*.

**Table S1. List of genes in the NMF7 signature.**

| Gene | | | | | | | | | | | |
|---|---|---|---|---|---|---|---|---|---|---|---|
| 1700025G04Rik | Atf2 | Cnot1 | Etl4 | Hdac8 | Man1a2 | Nek7 | Phactr4 | Rap1gds1 | Slc44a1 | Tbl1xr1 | Utrn |
| 1810026B05Rik | Atf7 | Cnot2 | Etv6 | Hectd1 | Man2a1 | Nemf | Phf14 | Rapgef5 | Slk | Tcf12 | Vps13a |
| 2610035D17Rik | Atf7ip | Cnot6l | Evi5 | Helz | Map2k4 | Nf1 | Phf20l1 | Rapgef6 | Slmap | Tcf20 | Vps13b |
| 2610307P16Rik | Atp11b | Col4a3bp | Ewsr1 | Herc1 | Map4k5 | Nfat5 | Phf21a | Rasa1 | Smarca2 | Tcf7l2 | Vps37a |
| 4632427E13Rik | Atp2b1 | Col4a5 | Exoc4 | Hivep2 | Mapkap1 | Nfia | Phip | Rasal2 | Smarcad1 | Tent2 | Vps54 |
| 4833423E24Rik | Atp6v1h | Cop1 | Exoc6b | Hook1 | Mark3 | Nfib | Phkb | Rbbp8 | Smc5 | Thoc2 | Vti1a |
| 4930402H24Rik | Atrnl1 | Copg2 | Ext1 | Hook3 | Mast2 | Nfkb1 | Phlpp1 | Rbm25 | Smg1 | Tia1 | Wac |
| 5031425E22Rik | Atrx | Cpsf6 | Faf1 | Huwe1 | Mbd5 | Nhlrc2 | Pik3c2a | Rbm39 | Smg6 | Tial1 | Wdfy3 |
| 9930021J03Rik | Atxn1 | Crebbp | Fam126b | Igf1r | Mbtd1 | Nipal2 | Pik3cb | Rbm5 | Smurf2 | Tlk1 | Wnk1 |
| Abi1 | Atxn2 | Cspp1 | Fam135a | Ikzf2 | Mcc | Nipbl | Pip4p2 | Rbms1 | Smyd3 | Tlk2 | Wwox |
| Adgrg6 | Babam2 | Ctnnd1 | Fam172a | Immp2l | Mdm4 | Nlk | Pitpnc1 | Rbms3 | Snx13 | Tmcc1 | Wwp1 |
| Adgrl2 | Baz1a | Ctnnd2 | Fam193a | Itch | Med13 | Npc1 | Pkp4 | Rcor1 | Sorbs2 | Tnks | Xpo7 |
| Afdn | Baz2b | Cux1 | Fam49b | Itfg1 | Med13l | Npepps | Plekha1 | Rel | Sox6 | Tnpo1 | Yes1 |
| Aff1 | Bbx | Dcaf8 | Far1 | Itga2 | Med14 | Nr6a1 | Plxdc2 | Rere | Spag9 | Tnrc6a | Ylpm1 |
| Agps | Bcas3 | Ddhd1 | Fbxl17 | Itsn2 | Mef2a | Nrd1 | Pnisr | Rfx7 | Specc1 | Tnrc6b | Ythdc1 |
| Ahcyl1 | Bdp1 | Ddi2 | Fbxl5 | Jmjd1c | Memo1 | Nrip1 | Pnn | Rictor | Spop | Tnrc6c | Ythdf3 |
| Aig1 | Birc6 | Ddx17 | Fbxo11 | Kansl1 | Met | Nsd1 | Ppm1a | Rin2 | Spred1 | Tns3 | Zc3h14 |
| Airn | Braf | Ddx5 | Fbxw7 | Katnbl1 | Mical3 | Nsd3 | Ppp1r12a | Rlf | Spred2 | Togaram1 | Zc3h7a |
| Akap13 | Brwd3 | Ddx50 | Fcho2 | Kdm6a | Mier1 | Nt5c2 | Ppp2r3a | Rnf19a | Sptlc2 | Top1 | Zcchc7 |
| Akt3 | Btbd7 | Dennd1a | Fchsd2 | Kidins220 | Mir205hg | Numa1 | Ppp2r5e | Rnf216 | Srgap2 | Tra2a | Zfand3 |
| Alcam | Camk1d | Dennd1b | Fer | Kif13a | Mkln1 | Numb | Ppp3ca | Robo1 | Srpk2 | Trim33 | Zfand5 |
| Ambra1 | Camsap2 | Dennd4c | Fgfr2 | Kif1b | Mllt10 | Nup98 | Ppp4r3b | Robo2 | Srrm2 | Trim44 | Zfc3h1 |
| Ank3 | Carmil1 | Diaph2 | Fip1l1 | Klhl24 | Mllt3 | Oga | Ppp6r3 | Rps6ka3 | Srsf11 | Trip12 | Zfp148 |
| Ankhd1 | Cask | Dip2b | Fnip1 | Kmt2c | Mob3b | Ogt | Prkca | Rps6kb1 | Srsf5 | Trpm7 | Zfp207 |
| Ankib1 | Cbfb | Dlg1 | Frk | Kmt2e | Mpp5 | Ophn1 | Prpf39 | Rsbn1l | Ssh2 | Trps1 | Zfp280d |
| Ankrd10 | Cblb | Dmxl1 | Fryl | Krit1 | Mpp7 | Osbpl3 | Prpf4b | Rsf1 | St5 | Tshz2 | Zfp292 |
| Ankrd11 | Ccnl1 | Dnajc1 | Fubp1 | Large1 | Mrtfa | Osbpl9 | Psme4 | Rsrc1 | Stag1 | Tshz3 | Zfp462 |
| Ankrd17 | Ccny | Dock1 | Fut9 | Larp4b | Msi2 | Oxr1 | Ptbp2 | Rtn4 | Stag2 | Ttc39b | Zfp53 |
| Aopep | Cdc73 | Dock11 | Fxr1 | Lars2 | mt-Atp8 | Pak1 | Pten | Runx1 | Stim1 | Tut4 | Zfp608 |
| Ap3b1 | Cdk12 | Dock7 | Galnt18 | Lcor | mt-Nd2 | Pak3 | Ptk2 | Samd12 | Stk3 | Ube2h | Zfp644 |
| Arap2 | Cdk13 | Dpyd | Gbf1 | Lcorl | mt-Nd4l | Pan3 | Ptpn13 | Sbf2 | Stox2 | Ube3c | Zfr |
| Arfgef1 | Cdk17 | Dst | Glg1 | Ldlrad4 | mt-Nd5 | Pard3 | Ptprj | Scaf8 | Strn3 | Ubn2 | Zfx |
| Arglu1 | Cdk19 | Ecpas | Gls | Lgr4 | Mtss1 | Parm1 | Pum1 | Scaper | Stx6 | Ubr3 | Zkscan3 |
| Arhgap10 | Cdk8 | Efna5 | Gm12602 | Lncpint | Mycbp2 | Parp8 | Pum2 | Scmh1 | Stx7 | Ubr5 | Zmym2 |
| Arhgap29 | Cers6 | Ehbp1 | Gm29904 | Lpin2 | Myo1b | Patj | Qk | Sema3d | Supt20 | Ufl1 | Zmym4 |
| Arhgap32 | Chd2 | Eif4a2 | Gm49890 | Lrba | Myo1d | Pawr | R3hdm1 | Senp6 | Susd6 | Uhrf2 | Znrf1 |
| Arhgap5 | Chd6 | Eif4g3 | Gmds | Lrrk1 | Myo1e | Pbx1 | R3hdm2 | Setd5 | Syne2 | Usp15 | |
| Arhgef12 | Chd9 | Ell2 | Gnaq | Luc7l2 | Myo6 | Pcm1 | Rab11fip2 | Sgms1 | Taf15 | Usp24 | |
| Arhgef3 | Chfr | Enah | Gpatch8 | Luzp1 | Naaladl2 | Pde4d | Rabep1 | Sh3bgrl | Tanc1 | Usp25 | |
| Arid4a | Chka | Ep300 | Gpbp1l1 | Macc1 | Nav2 | Pde7a | Rabgap1 | Sh3d19 | Tanc2 | Usp3 | |
| Arid4b | Clcn3 | Epha7 | Gphn | Macf1 | Nckap1 | Pdlim5 | Rabgap1l | Sh3kbp1 | Taok1 | Usp32 | |
| Arl15 | Clk1 | Eps15 | Grip1 | Magi1 | Ncor1 | Pds5a | Rad51b | Sh3rf1 | Taok3 | Usp34 | |
| Asap1 | Clock | Erbin | Grm8 | Magi3 | Neat1 | Pds5b | Ralgapa1 | Sipa1l1 | Tbc1d1 | Usp53 | |
| Asxl2 | Cmip | Erc1 | Gsap | Maml2 | Nebl | Peli1 | Ralgapa2 | Skap2 | Tbc1d5 | Usp6nl | |
| Atad2b | Cmss1 | Esyt2 | Gtf2h1 | Maml3 | Nedd9 | Phactr2 | Ranbp9 | Slc24a3 | Tbl1x | Usp9x | |
