## Supplementary Figures for "Deciphering Precursor Cell Dynamics in Esophageal Preneoplasia via Genetic Barcoding and Single-Cell Transcriptomics"

### Supplementary Figure 1

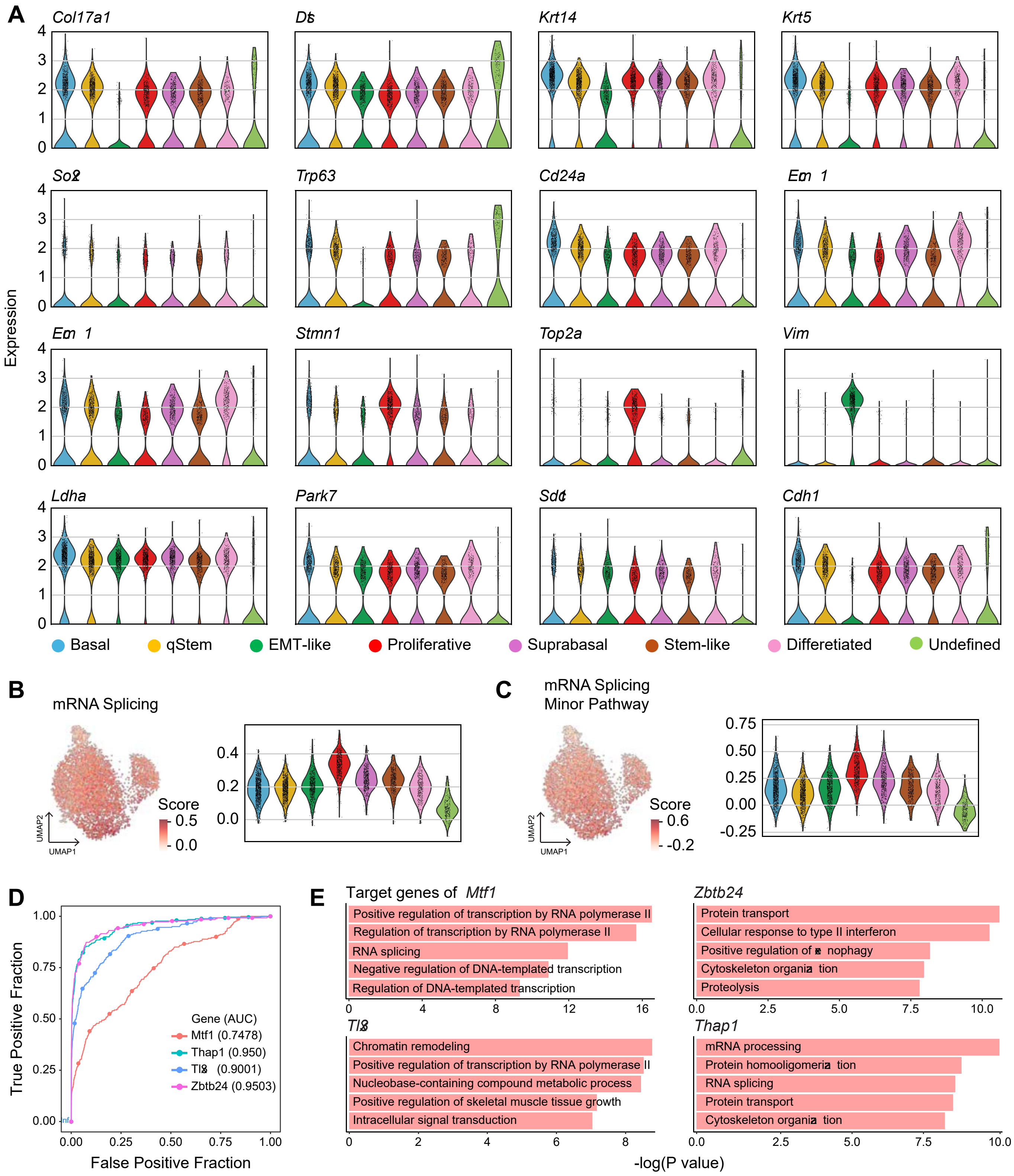

#### Supplementary Figure 2

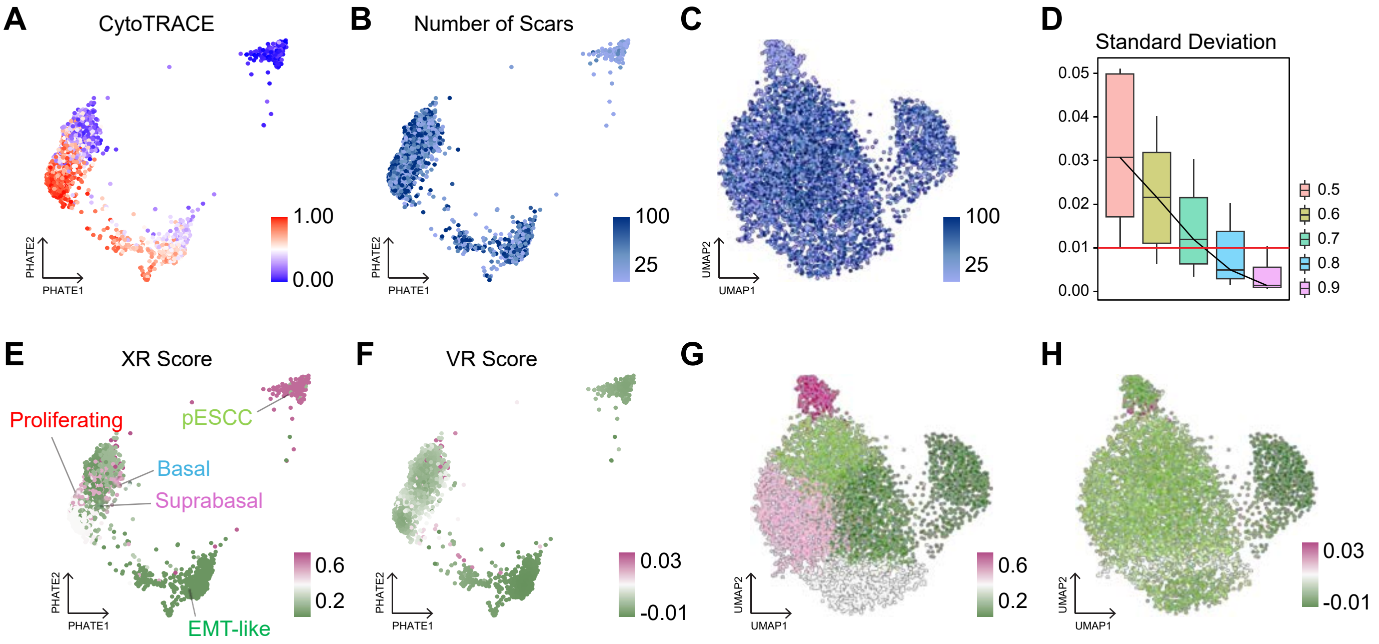

Supplementary Figure 3

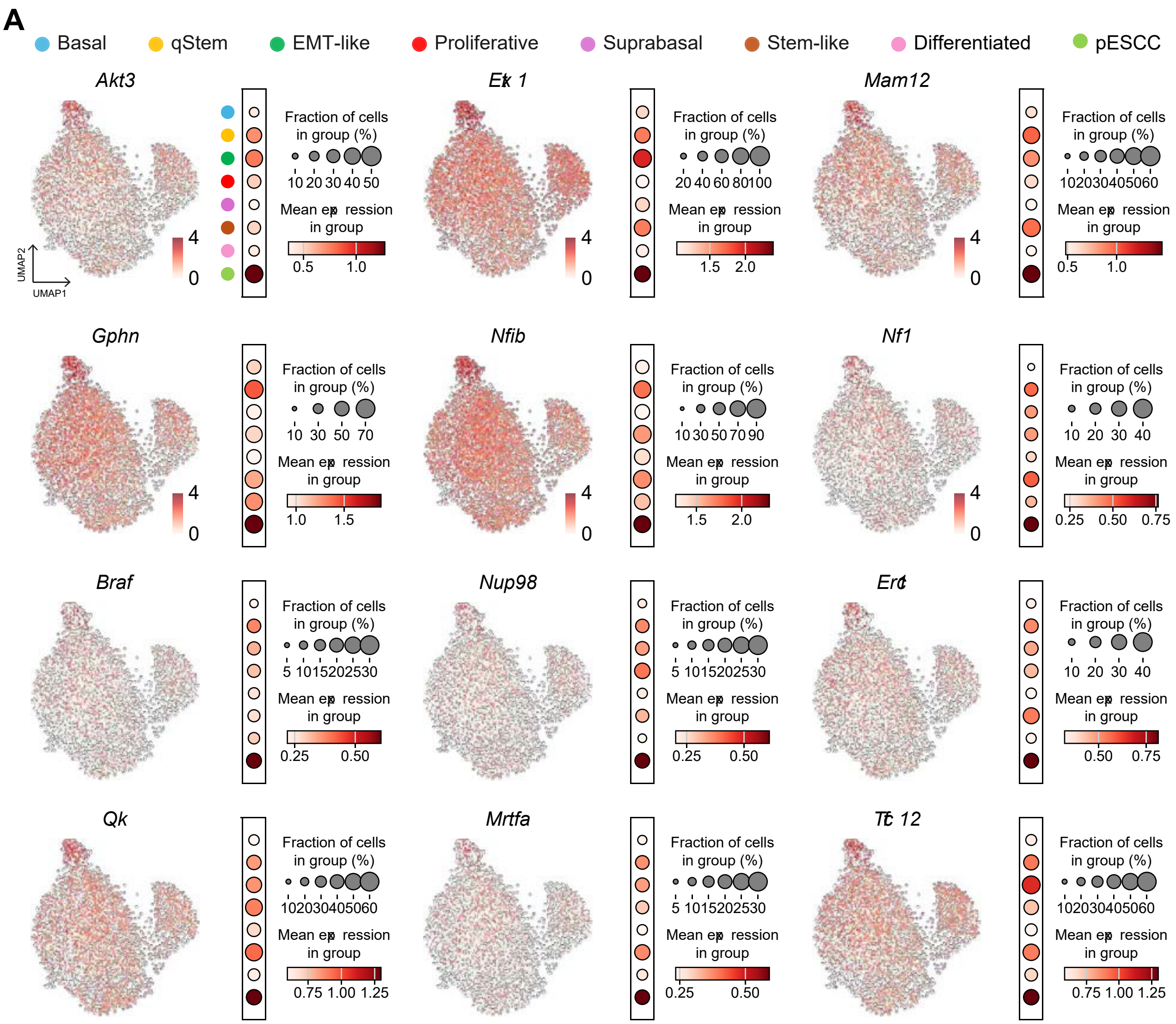

### Supplementary Figure 4

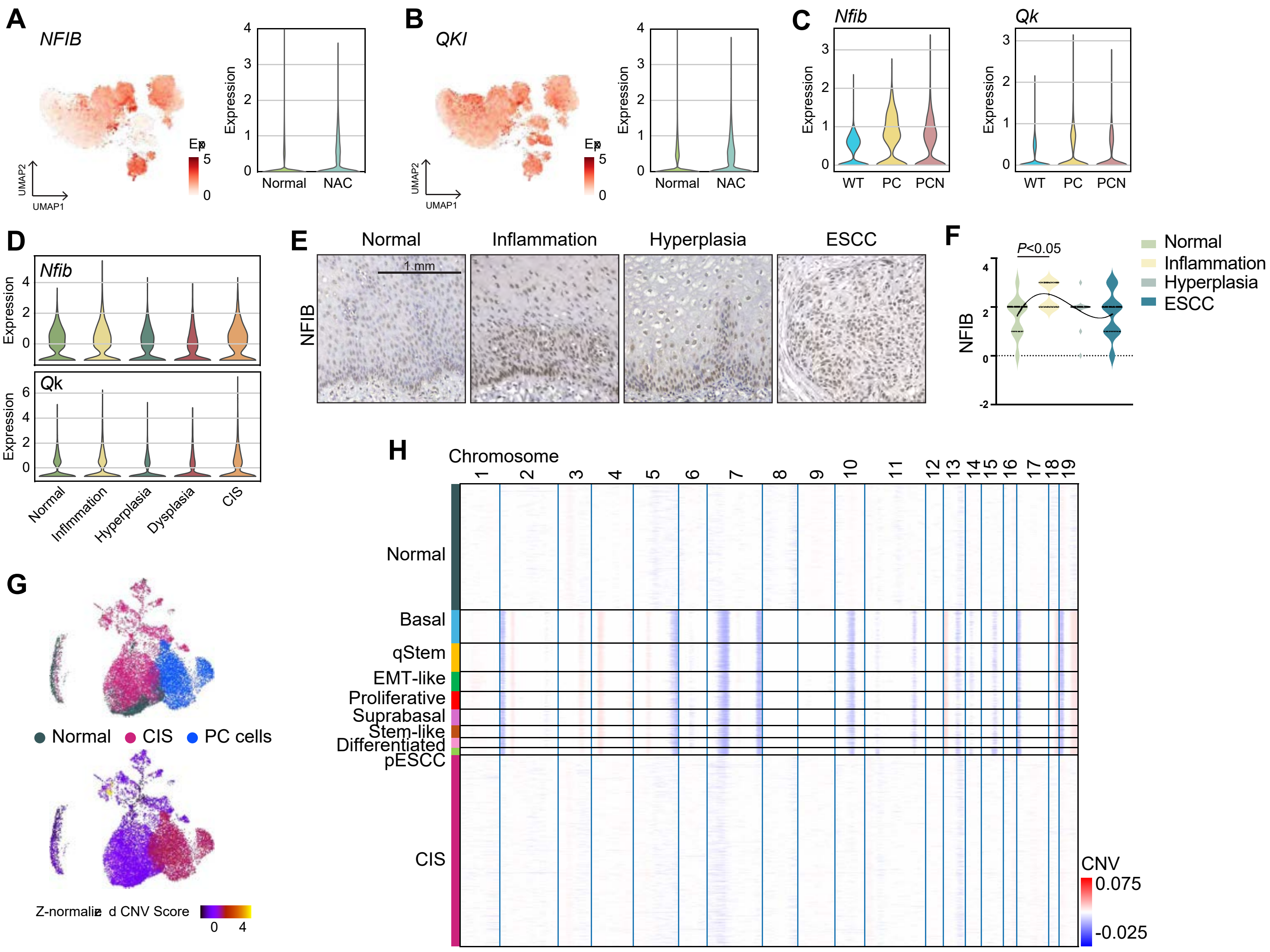

#### Supplementary Figure 5

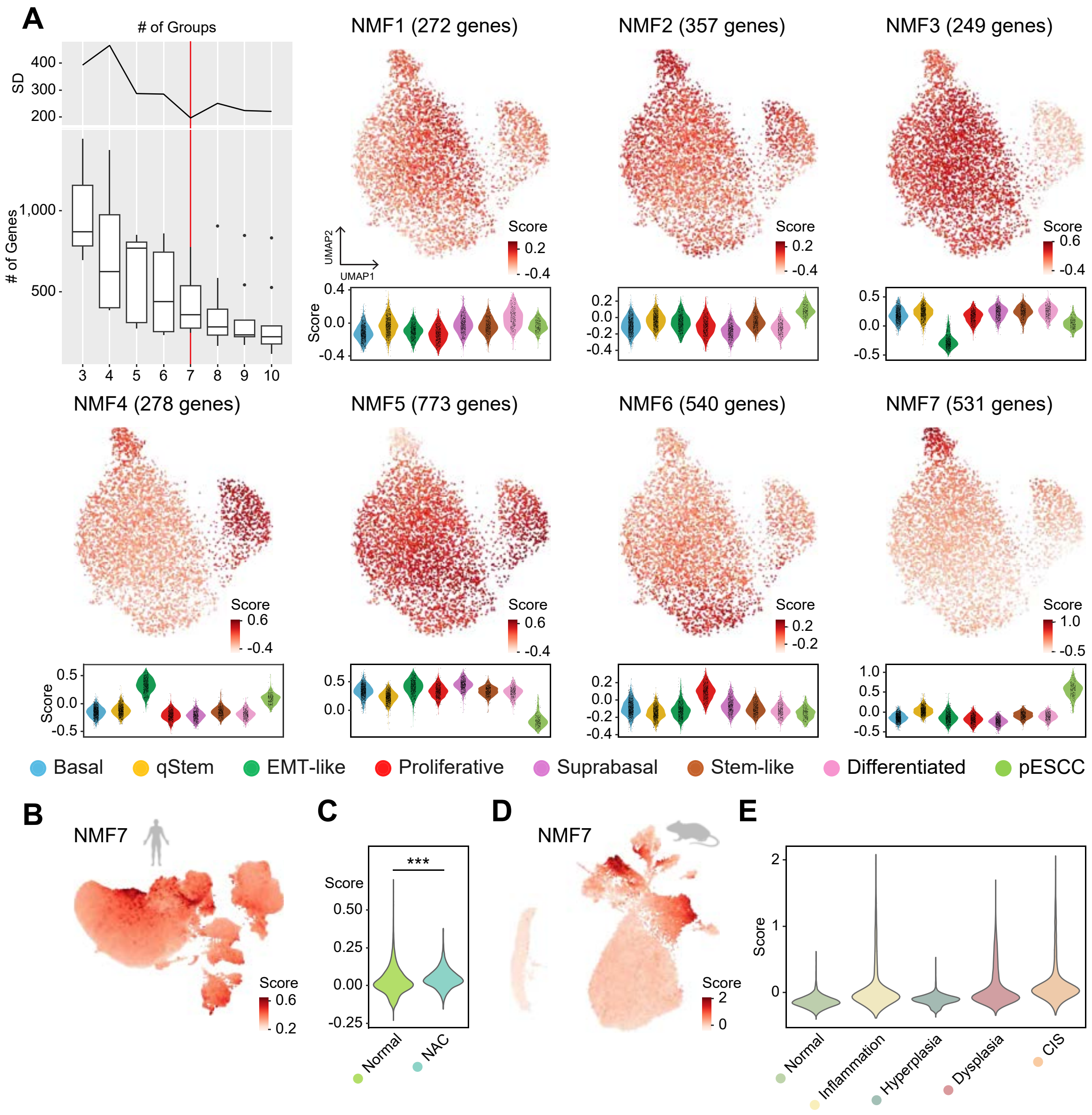
